# Interplay between somatic and dendritic inhibition promotes the emergence and stabilization of place fields

**DOI:** 10.1101/483875

**Authors:** Victor Pedrosa, Claudia Clopath

## Abstract

During exploration of novel environments, place fields are rapidly formed in hippocampal CA1 neurons. Place cell firing rate increases in early stages of exploration of novel environments but returns to baseline levels in familiar environments. However, although similar in amplitude and width, place fields in familiar environments are more stable than in novel environments. We propose a computational model of the hippocampal CA1 network, which describes the formation, the dynamics and the stabilization of place fields. We show that although somatic disinhibition is sufficient to form place cells, dendritic inhibition along with synaptic plasticity is necessary for stabilization. Our model suggests that place cell stability is due to large excitatory synaptic weights and large dendritic inhibition. We show that the interplay between somatic and dendritic inhibition balances the increased excitatory weights, so that place cells return to their baseline firing rate after exploration. Our model suggests that different types of interneurons are essential to unravel the mechanisms underlying place field plasticity. Finally, we predict that artificial induced dendritic events can shift place fields even after place field stabilization.

## 1 Introduction

The hippocampus encodes spatial representations. A subset of hippocampal pyramidal cells— called place cells—fire action potentials when the animal is in a specific location within the environment, the place fields [O’Keefe and Dostrovsky, 1971, O’Keefe, 1976, O’keefe and Nadel, 1978, Wilson and McNaughton, 1993]. How these place fields are formed is not clear yet. In particular, experimental data open up puzzling questions:

Subthreshold responses of silent cells, when recorded at the soma, are not place-tuned [Epsztein et al., 2011]. If a spatially uniform current is applied to silent cells, however, these cells start to produce place-tuned activity [Lee et al., 2012]. This transition from silent to place cell is abrupt. This result suggests that silent cells receive place-tuned inputs but there is no signature of those inputs at the soma. **How can silent cells become place cell despite the fact that their subthreshold response is untuned?**

There is increasing evidence suggesting that place fields are not formed from scratch [Cacucci et al., 2007, Dragoi and Tonegawa, 2011, Dragoi and Tonegawa, 2013, Dragoi and Tonegawa, 2014, Frank et al., 2004, Hill, 1978, Kentros et al., 1998, McHugh et al., 1996, Sheffield et al., 2017, Epsztein et al., 2011]. Instead, spatial representation might be built from the selection of already strong connections, without the need for synaptic plasticity [Dragoi and Tonegawa, 2011, Dragoi and Tonegawa, 2013, Dragoi and Tonegawa, 2014, Lee et al., 2012]. For instance, many place fields, although not stable, are present since the’s first exploration of a novel environment [Epsztein et al., 2011, Frank et al., 2004, Hill, 1978]. **How can CA1 pyramidal cells become place cells without synaptic plasticity?**

During exploration of a novel linear track, new place fields are formed over several laps [Sheffield et al., 2017]. The development of these new place fields have been shown to be preceded by dendritic regenerative events—back propagating action potentials or dendritically generated spikes [Sheffield et al., 2017, Sheffield and Dombeck, 2015]. Moreover, exposure to novel environment is associated with a reduction in dendritic inhibition. More recently, place fields have been produced by the conjunctive activation of presynaptic inputs and postsynaptic calcium plateau potentials [Bittner et al., 2017]. Place fields have also been developed following juxtacellular stimulation of CA1 silent cells [Diamantaki et al., 2018]. **What is the relationship between dendritic activity and place field formation?**

Place cell firing rate is initially low but increases rapidly during exploration of novel environments [Cohen et al., 2017]. Surprisingly, in familiar environments, place cell activity returns to a lower level, comparable to the activity during initial exploration of novel environments [Cohen et al., 2017]. **How can place cell activity return to baseline firing rate after a transient increase in firing at the beginning of exploration?**

Remarkably, although place cell activity is similar in the first stages of exploration of novel environments and in familiar environments, place fields in familiar environments have been shown to be considerably more stable [Cohen et al., 2017]. Synaptic plasticity has been shown in be involved in this stabilization [Cohen et al., 2017]. Moreover, the blockage of NMDA receptors in CA1 neurons has been shown to significantly decrease the number of place fields formed across the network. These results suggest that synaptic plasticity is not required for the formation of place fields but is involved in the development of new place cells and stabilization of spatial representations. **Despite the fact that place cells in novel and familiar environments have similar activity, how can they have different levels of stability?**

To address these questions, we develop a data-driven model of the hippocampal CA1 network. We show that somatic disinhibition is sufficient to form place cells. However, dendritic inhibition and synaptic plasticity allows for silent cells to turn into stable place cells. We show that the combined action of somatic and dendritic inhibition balances an increase in excitatory weights due to synaptic plasticity, so that place cells after exploration return to their baseline firing rate. Our model suggests that place cell stability is due to large excitatory synaptic weights and large dendritic inhibition. Therefore, our model suggests that different types of interneurons are essential to unravel the mechanisms underlying place field plasticity. Finally, we use our model to predict how to perturb place fields. Artificially induced dendritic events in place cells can shift place field location even after place field stabilization. Our model reproduces a wide range of observations from the hippocampal CA1 network, provides a circuit level understanding, and finally makes predictions that can be tested in future experiments. Importantly, our model suggests that interneuron diversity is crucial for the emergence of place fields and their consolidation.

## 2 Results

In all simulations, we model CA1 pyramidal neurons as two-compartment, rate-based neurons (figure 1). The neurons have non-linear dendritic units to account for dendritic spikes (figure 1). We assume that place-tuned inputs are projected onto dendrites of all CA1 cells and the propagation of inputs from dendrites to soma is gated by somatic depolarization (figure 1). For the sake of simplicity, synaptic plasticity depends on presynaptic activity and the postsynaptic dendritic activation only (figure 1, see methods for details). Finally, exploration of novel environments has been shown to modulate CA1 interneuron activity in an interneuron-type-specific manner [Sheffield et al., 2017]. In our model, we hypothesize the existence of a novelty signal and we assume that interneuron activity is modulated by this novelty signal (figure 1, see methods). In particular, dendrite-targeting inhibition is decreased in novel environments and slowing returns to baseline. Conversely, soma-targeting inhibition is increased in novel environments followed by a slow decay to baseline.

**Figure 1.**
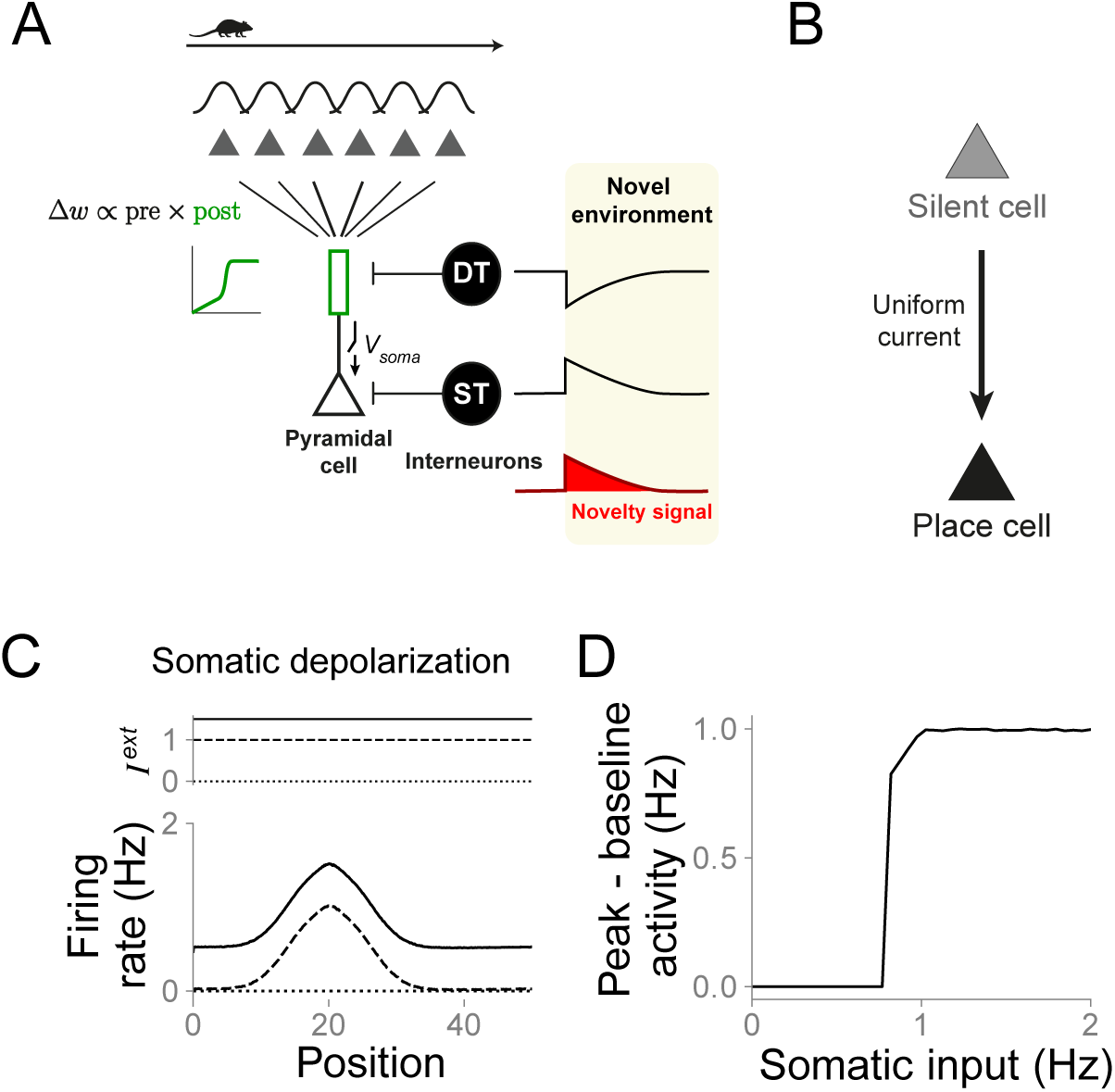
Somatic disinhibition is sufficient to turn silents cell into place cells. **(A)** Network diagram. Pyramidal neurons receive place-tuned, excitatory input and inputs from two types of interneurons: dendrite-targeting (DT), representing somatostatin-expressing interneurons, and soma-targeting (ST), representing parvalbumin-expressing interneurons. The activity of interneurons is modulated during the exploration of novel environments. DT interneuron activity (top black curve) decreases, whereas ST interneuron activity (bottom black curve) increases in novel environments. Both interneuron activities gradually return to baseline levels with a timescale defined by the hypothesized novelty signal (red curve, see methods and main text for details). **(B)** Diagram of a silent cell being turned into a place cell following uniform depolarization. **(C)** Pyramidal cell firing rate as a function of the animal position for three different levels of depolarization. Top, external current applied to the somatic compartment. Bottom, pyramidal cell firing rate for the three corresponding external currents. **(D)** Difference between peak and baseline firing rate as a function of the external somatic input. Because of the gated propagation of inputs from dendrites to soma, there is an abrupt transition from silent to place cell.

### 2.1 Somatic disinhibition is sufficient to turn silents cell into place cells

We first investigate how silent cells can be transiently turned into place cells through the injection of a spatially uniform current. We simulate 10 input neurons, which could be thought of as part of CA3, projecting onto one postsynaptic CA1 neuron. The CA1 neuron also receives dendritic- projecting inhibition (thought as a subset of somatostatin-expressing cells) and somatic-projecting inhibition (thought as a subset of parvalbumin-expressing cells). For simplicity, all presynaptic neurons are assumed to have constant, uniformly distributed place fields. We assume that the postsynaptic neuron receives tuned input from input neurons. Although not uniform, the initial synaptic weights are chosen such that the postsynaptic CA1 cell is silent during the first lap of exploration. During the second lap of exploration, an external depolarizing current is applied to the somatic compartment. Since plasticity is slow, synaptic weights are not significantly changed from the first to the second lap. However, because the propagation of inputs from dendrites to soma is gated by somatic depolarization, silent cells are turned into place cell in an all-or-nothing manner (figure 1B-D). For weak external currents, silent cells remain silent. For sufficiently strong external currents, silent cells become place cells (figure 1C). Furthermore, the transition from silent to place cell is abrupt (figure 1D). Therefore, our model indicates that silent cells can be transiently turned into place cells due to a combination of two features: silent cells receive place-tuned input and the propagation of these inputs from dendrites to soma is gated by somatic depolarization.

### 2.2 Dendritic disinhibition and synaptic plasticity allows silent cells to turn into stable place cells

Using our model, we next investigate whether there is an alternative mechanism underlying place field formation of originally silent cells. As before, we simulate 10 input neurons projecting onto one postsynaptic CA1 neuron. Synaptic connections from input neurons to the postsynaptic neuron are plastic and their change depends on the activity of the postsynaptic dendritic compartment. Inspired by the experiments from Sheffield et al. [Sheffield et al., 2017], we simulate an “novelty signal”, so that the activity of dendrite-targeting interneurons is initially low in novel environments and increases gradually while the environment is getting familiar, whereas the activity of soma-targeting interneurons is initially high and decreases gradually (figure 2A-B). Simulating our model reveals that the reduction in dendrite-targeting inhibition opens the possibility for dendritic activity in the pyramidal cells, regardless of the somatic activity (figure 2C). This activation promotes the strengthening of synaptic weights (figure 2D-F). The combination of stronger synaptic weights and the decrease in soma-targeting inhibition finally leads to the development of place-tuned somatic activity (figure 2C). Therefore, our model suggests that the combination of dendritic-activity-dependent synaptic plasticity and novelty-modulated interneuron activity can turn silent cells into place cells. Interestingly, dendritic activity in simulated CA1 neurons precedes and predicts place field development in silent cells in our model, consistent with the experimental findings of Sheffield et al. [Sheffield et al., 2017].

**Figure 2.**
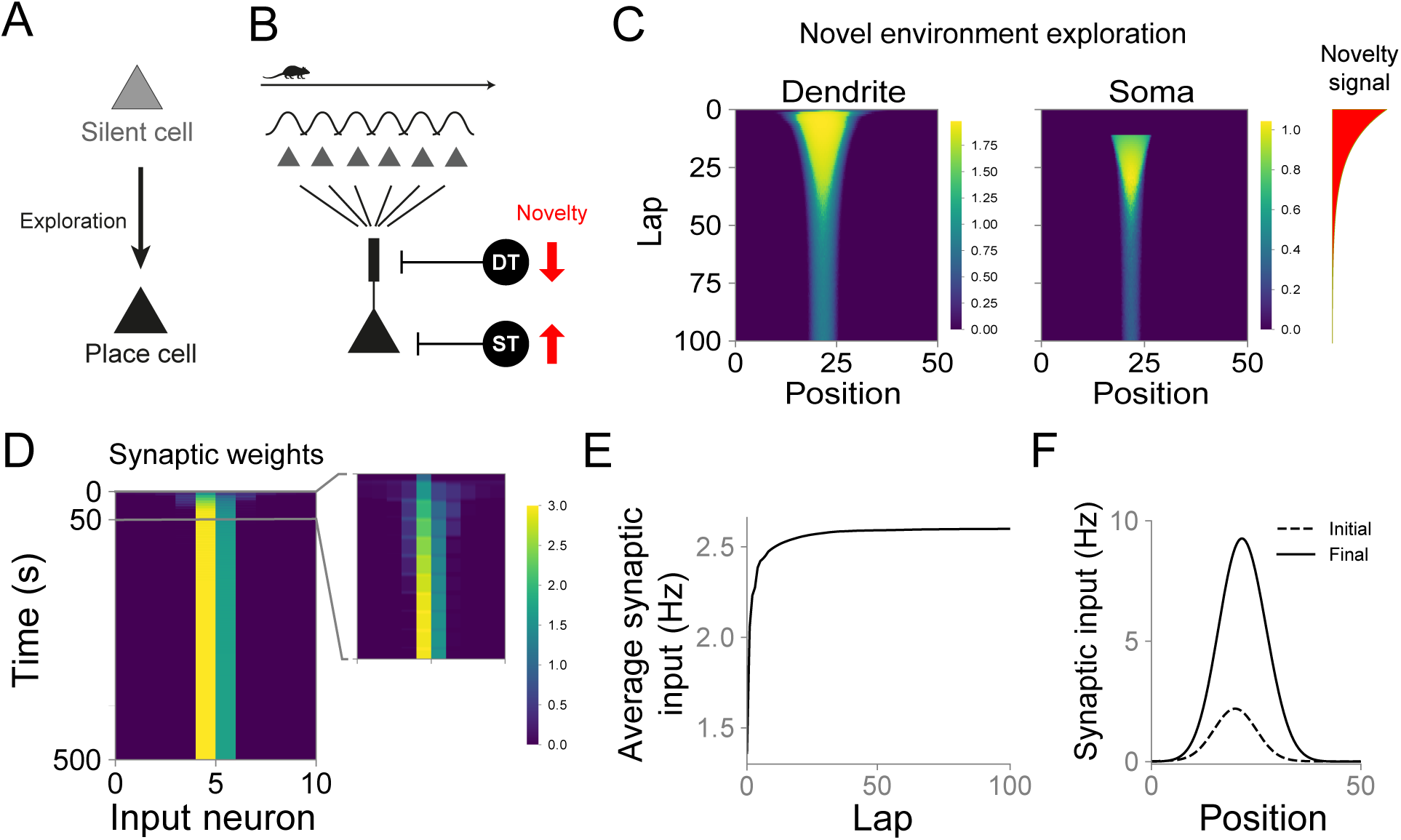
Dendritic disinhibition and synaptic plasticity allows silent cells to turn into stable place cells. **(A)** Diagram of a silent cell being turned into a place cell following the exploration of a novel environment. **(B)** Single-cell diagram. One postsynaptic neuron receives input from 10 presynaptic neurons whose activities are modulated by the animal’s position. The postsynaptic neuron is simulated as a two-compartment neuron. The neuron receives dendritic and somatic inhibition and the inhibitory input is modulated by novelty (see methods). **(C)** Silent cell turns into place cell following exploration of a novel environment. Evolution of dendritic (left) and somatic (middle) activity during exploration of a novel environment for an initially silent cell. Amplitude of novelty signal over laps (right, red). Dendritic activity precedes somatic activation, in agreement with experiments [Sheffield et al., 2017]. Somatic activity increases abruptly due to the gated propagation of dendritic inputs (see methods). **(D)** Evolution of synaptic weights for the same example cell shown in C. Inset: first 10% (50 s) of exploration. **(E)** Evolution of average synaptic input over laps for the same example cell as in C. **(F)** Initial (dashed) and final (solid) synaptic inputs as a function of the animal position for the same example cell as in C. The synaptic input was measured as the convolution between initial/final synaptic weights and the input neuron activities.

### 2.3 Interplay between somatic and dendritic inhibition balances increased excitatory synaptic weights so that place cells firing rate returns to baseline

We then study neurons in our model that are already place cells in a novel environment. As before, our model consists of a CA1 cell receiving place-tuned inputs. But here, the synaptic weights are such that the neuron is active since the first lap (figure 3A-D, blue traces). The initial low dendritic inhibition in our model leads to the activation of dendritic compartments and thus the strengthening of synaptic weights. Stronger synaptic weights produce a stronger neuronal response (figure 3A-D, green traces). However, when the environment starts to become familiar, the novelty signal vanishes.

**Figure 3.**
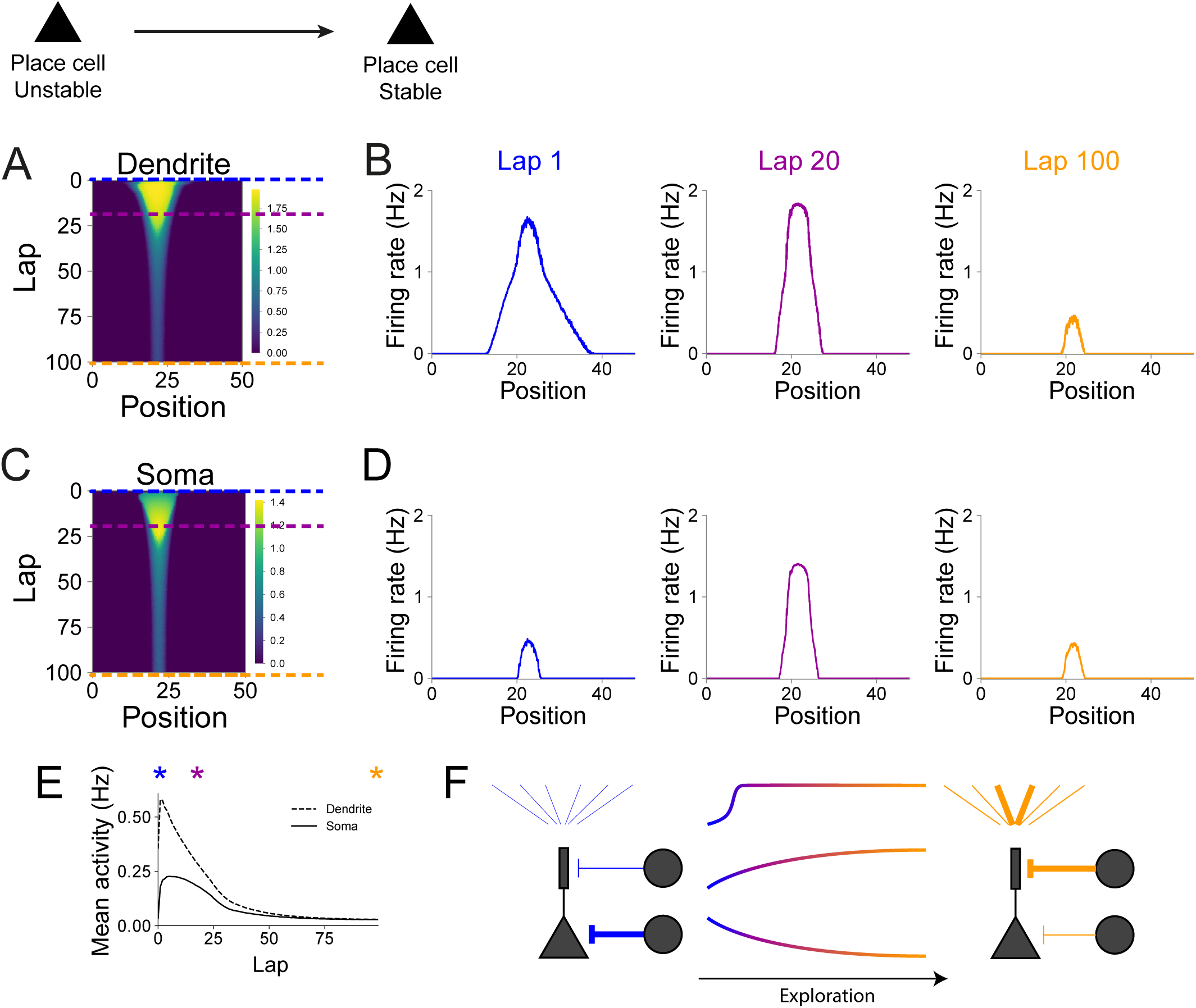
Interplay between somatic and dendritic inhibition balances increased excitatory synaptic weights so that place cells firing rate returns to baseline. **(A-H)** Place field dynamics for an initially active cell. **(A)** Evolution of dendritic activity for an example place cell. **(B)** Dendritic activity as a function of the animal’s position for three stages of the simulation: lap 1 (left, blue; blue dashed line in (A)), lap 20 (middle, dark green; dark green dashed line in (A)), and lap 100 (right, orange; orange dashed line in (A)). **(C)** Evolution of somatic activity for the same cell as in (A). **(D)** Somatic activity as a function of the animal’s position for three stages of the simulation: lap 1 (left, blue; blue dashed line in (C)), lap 20 (middle, dark green; dark green dashed line in (C)), and lap 100 (right, orange; orange dashed line in (C)). **(E)** Evolution of mean dendritic (dashed line) and somatic (solid line) activity for the same example cell as in (A) and (C). Stars indicate laps 1 (blue), 20 (dark green) and 100 (orange). Both somatic and dendritic activities increase sharply during the first laps of exploration due to synaptic plasticity. **(F)** Diagram showing the changes in the network from the first to the last lap of exploration. Initially (left, blue), input synaptic weights are weak, dendritic inhibition is low and somatic inhibition is high. During the final lap (right, orange), some input synaptic weights are strong, dendritic inhibition is high and somatic inhibition is low. Therefore, although place field amplitude and width are the same in the first and last lap (D blue and orange), the network is in a different state.

Thus the dendritic inhibition is higher, resulting in a lower activation of the dendritic compartment (figure 3B, orange trace). The lower level of somatic inhibition allows the neuron to exhibit the same level of activity even under reduced dendritic activation (figure 3D, orange trace, and figure 3E). Our model is therefore consistent with the experimental data showing that place fields gradually increase during the exploration of novel environments and later return to the initial levels in familiar environments [Cohen et al., 2017]. Importantly, although the neuronal firing on the first and last laps are indistinguishable, the network states are distinct. During the first lap, the neuron receives weak excitatory input and dendritic inhibition, and strong somatic inhibition. During the last lap (when the environment is familiar), the neuron receives strong excitatory input and dendritic inhibition, and weak somatic inhibition (figure 3F).

### 2.4 Large synaptic weights and large dendritic inhibition provide place cell stability

We next investigate whether place fields in familiar environments are more stable than at the beginning of the exploration phase in novel environments—despite both having the same amplitude and tuning width. In order to do that, we assume that the place field can be affected by three sources of noise: (i) noise on the place fields of presynaptic neurons, (ii) noise on the firing rates of presynaptic neurons, or (iii) noise on synaptic weights, accounting e.g. for synaptic turnover or synaptic failure (figure 4A). In all three cases, we compare the effect of noise on place fields at the beginning of exploration (figure 4, blue curves) to its effect on place fields at the end of exploration (figure 4, orange curves; see methods). In case (i), we assume that the amplitudes of presynaptic place fields are not all the same. Instead, we multiply each place field by a random number whose variance increases with the noise amplitude (see methods). Of course, the more noise we impose, the less stable place cells are (figure 4A). However, the noise on presynaptic place fields is more effective at destabilizing place cells in the first lap of exploration compared to at the end of exploration (figure 4A), suggesting that place cells become more stable. In case (ii), we assume that all presynaptic place fields have the same amplitude but input neurons can also fire at any time with probability *p*. This probability increases linearly with noise amplitude. Again, place fields at the final lap are more stable than initial place fields (figure 4B). In case (iii), we change synaptic weights by random amounts whose variance is proportional to the noise amplitude. This source of noise also affects initial place fields more than it does to final place fields (figure 4C). In all three cases, the stabilization of place fields results from increased synaptic weights and higher dendritic inhibition (figure 3F). Therefore, place fields in familiar environments are more stable to noise than place fields at the beginning of novel environment exploration, consistent with experimental observations [Cohen et al., 2017].

**Figure 4.**
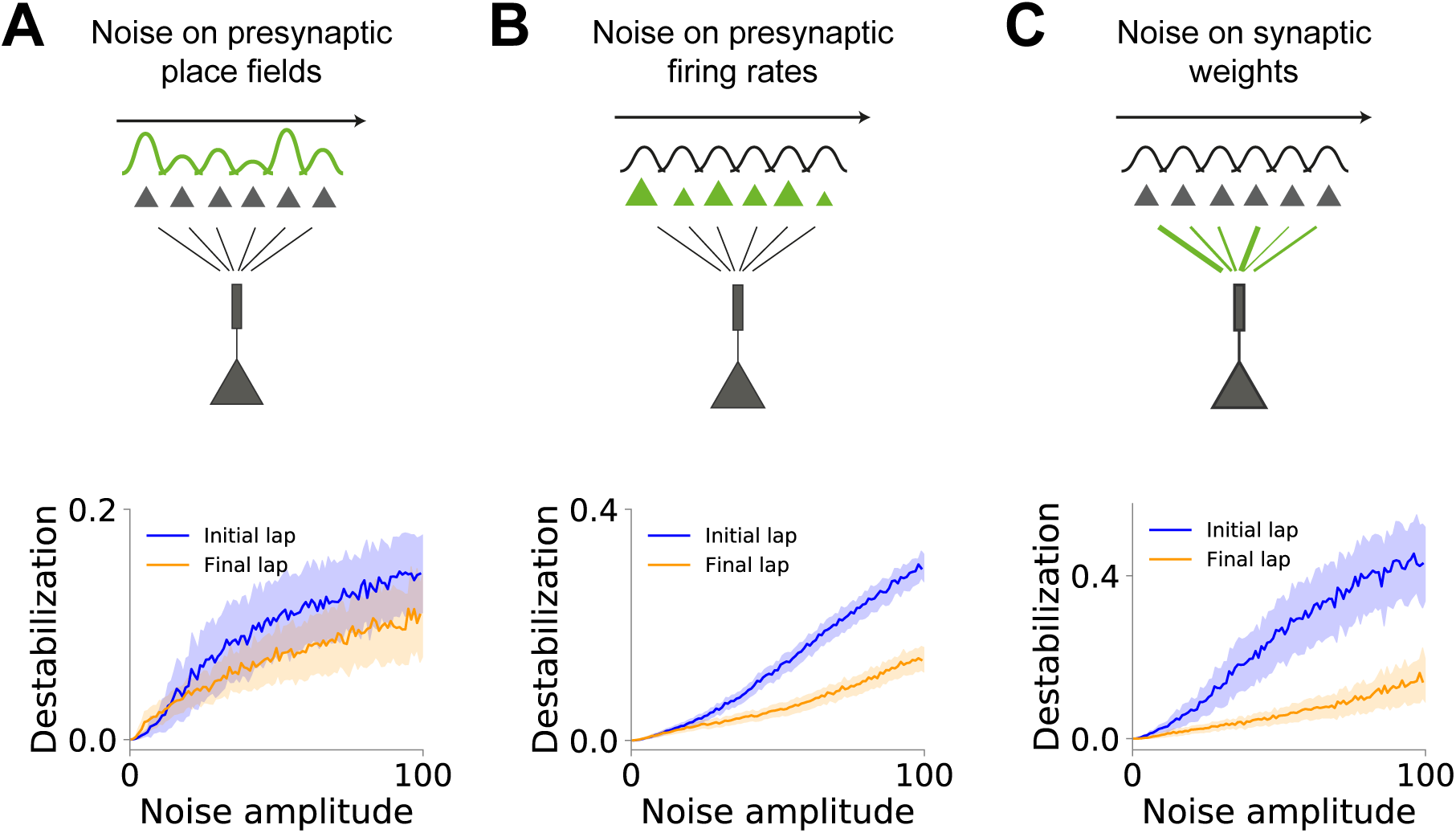
Large synaptic weights and large dendritic inhibition provide place cell stability. **(A-C)** Effect of noise on place fields for the first (blue) and last (orange) laps of exploration. **(A)** Destabilization of place fields by noise on presynaptic place fields. We measure the change on postsynaptic place field following changes on presynaptic place field amplitudes (see methods). **(B)** Destabilization of place fields by noise on presynaptic firing rates. We measure the change on postsynaptic place field following the addition of a noisy input to presynaptic neurons (see methods). **(C)** Destabilization of place fields by noise on synaptic weights. We measure the change in postsynaptic place field following changes on synaptic weights (see methods). For all three sources of noise (A-C), the effect of the noise over place fields is higher in the first lap than in the last lap.

In order to investigate the role of each component of the network in stabilizing place fields, we artificially modify the final state of the network while keeping the neuron’s place field unchanged. We first reduce the amplitude of both excitatory weights and dendritic inhibition (supplementary figure 1A). The reduced synaptic weights decrease place field stability when noise is added on the synaptic weights (supplementary figure 1B). Next, we reduce dendritic inhibition and increase somatic inhibitory input (supplementary figure 1C). Since synaptic weights are strong and dendritic inhibition is low, the postsynaptic neuron is more susceptible to presynaptic inputs. Thus, noise on presynaptic neurons is carried on to postsynaptic place fields, destabilizing them (supplementary figure 1D). In summary, strong synaptic connections are relatively less affected by noise on synaptic weights, whereas higher dendritic inhibition cancels out-of-field fluctuations being transmitted from presynaptic neurons.

We next investigate whether dendritic nonlinearity can contribute to stable place field development. In our model, when inputs are strong enough, they can induce dendritic spikes, which in turn lead to strong potentiation. As such, dendritic spikes—or dendritic nonlinearities—might form a mechanism for reliably selecting presynaptic inputs. To test this hypothesis, we simulate our model with initially uniform synaptic weights and no novelty signal. We then compare it with an alternative model where dendrites do not have a nonlinearity but can reach the same maximum level of activity (linear dendrites, supplementary figure 2A-B). Neurons with dendritic nonlinearity develop place fields faster and, importantly, more reliably (supplementary figure 2C). In several cases, neurons with linear dendrites do not develop place fields and their activity varies from lap to lap (supplementary figure 2D). Contrarily, neurons with dendritic nonlinearity consistently develop stable place fields. Therefore, our model suggests that dendritic nonlinearities might contribute to place field development and stability by promoting a reliable selection of inputs.

### 2.5 Artificially induced dendritic events induce place field plasticity

Using our model, we next explore whether it is possible to perturb or change single CA1 place fields. We simulate a single neuron receiving place-tuned input such that one of its input synapses is stronger than the remaining connections. We assume that the animal is exploring a novel environment. As such, interneuron activity is modulated by a novelty signal that decays over time (figure 5A, see methods). The stronger synaptic weight leads to the activation of our neuron, which leads to the strengthening of that synaptic weight. This positive feedback loop leads to the development of a strong place field (figure 5B).

**Figure 5.**
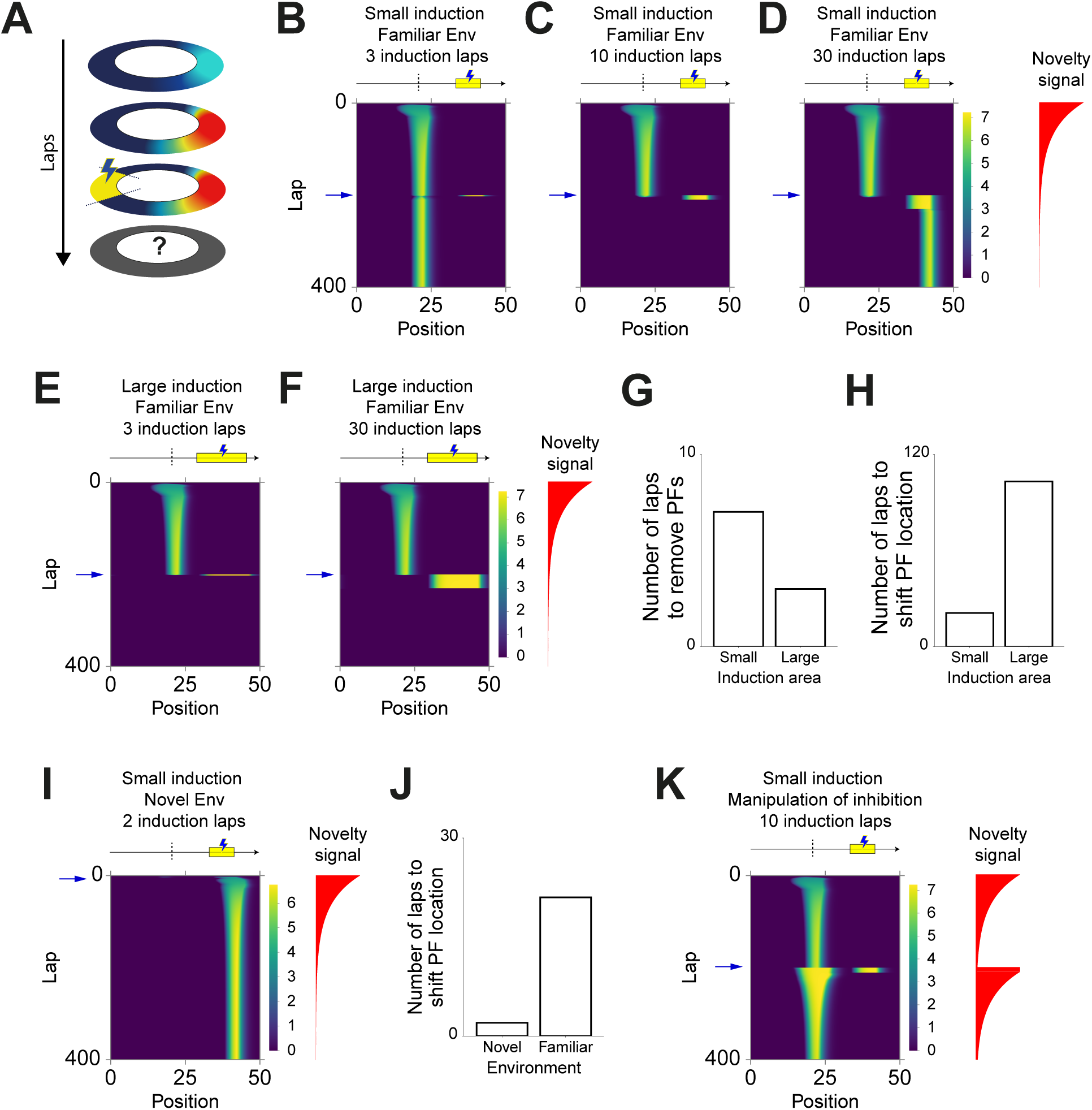
Artificially induced dendritic events induce place field plasticity. **(A)** Single-cell diagram. **(B-D)** Evolution of place fields for the case in which an extra current is applied to the postsynaptic neuron while the animal traverses a small section (1/6) of the track. Yellow bar indicates the induction region in which the extra current is applied. Dashed line indicates the position of the peak of the initial place field. Blue arrow indicates the first induction lap (lap 200). Red curve shows the evolution of the novelty signal over laps. **(B)** Place field evolution for 3 induction laps. Place fields are not disturbed following the application of extra current. **(C)** Place field evolution for 10 induction laps. Place fields are removed by the application of extra current. **(D)** Place field evolution for 30 induction laps. Place fields are shifted towards a new position determined by the region of extra current application. **(E-F)** Evolution of place fields for the case in which an extra current is applied to the postsynaptic neuron while the animal traverses a large section (2/6) of the track. Yellow bar indicates the induction region in which the extra current is applied. Dashed line indicates the position of the peak of the initial place field. Blue arrow indicates the first induction lap (lap 200). Red curve shows the evolution of the novelty signal over laps. **(E)** Place field evolution for 3 induction laps. Place fields are removed by the application of extra current. **(F)** Place field evolution for 30 induction laps. Place fields are removed by the application of extra current. **(G)** Number of induction laps required to remove stable place field for small and large induction areas. **(H)** Number of induction laps required to shift place field location for small and large induction areas. **(I)** Evolution of place fields for the case in which an extra current is applied during exploration of a novel environment (lap 2). The extra current is applied to the postsynaptic neuron while the animal traverses a small section (1/6) of the track. Yellow bar indicates the induction region in which the extra current is applied. Dashed line indicates the position of the peak of the initial place field. Blue arrow indicates the first induction lap (lap 2). Red curve shows the evolution of the novelty signal over laps. **(J)** Number of induction laps required to shift place field location for novel and familiar environments. **(K)** Evolution of place fields for the case in which the application of an extra current is paired with the resetting of the novelty signal. The extra current is applied to the postsynaptic neuron while the animal traverses a small section (1/6) of the track. Yellow bar indicates the induction region in which the extra current is applied. Dashed line indicates the position of the peak of the initial place field. Blue arrow indicates the first induction lap (lap 2). Red curve shows the evolution of the novelty signal over laps. The novelty signal resetting leads to a reduction in dendritic inhibition across the whole track. Therefore, the in-field activity increases, leading to the reinforcement of the initial place field.

We then test whether we can shift the tuning of the place field towards a new location by artificially activating CA1 neurons. In order to do that, we simulate the network until the novelty signal is negligible—the environment is hence considered familiar—and the postsynaptic place field is stable. At this stage, we inject an extra current in the dendritic compartment of the simulated neuron to induce a strong dendritic activity. This current is induced only in a small region within the track, far from the peak of the postsynaptic place field (figure 5, see methods). The induction of extra dendritic activity over a few (3) laps does not alter the postsynaptic place field (figure 5B). We next induce the extra dendritic activity over several (30) laps. In this case, the position of the place field is shifted towards the new location (figure 5D). For an intermediate number of induction laps, the initial place field is removed without the formation of a new place field, thus turning the place cell into a silent cell (figure 5C). Note that this newly formed silent cell can potentially develop a place field again, in the case where there is remaining dendritic activity. This dendritic activity allows for plasticity, and therefore for the re-emergence of a place field (supplementary figure 3). Altogether, our model predicts that, if induced over enough laps, artificial dendritic activity can shift place field location.

The size of the induction region might affect the efficacy to shift place field location. To investigate this, we increase the induction area to twice its original size. In this case, the induction over three laps is enough to remove the initial place field, but not enough to induce the formation of a new one (figure 5E). Induction over 30 laps—which is enough to induce the development of a new place field for a small induction area—is not enough to promote the development of a new place field (figure 5F). The larger the induction area, the easier it is to remove the initial place tuning (figure 5G). Nevertheless, a large induction area leads to a competition between inputs within that area. Because of that, our model predicts that, surprisingly, the larger the induction region, the more induction laps are needed to induce the development of new receptive fields (figure 5H).

We next compare the induction of place field shift in novel and familiar environments. We hy- pothesize that in novel environments, place fields should be more plastic and, therefore, it should be easier to induce a shift in place field location. In order to test this, we induce dendritic activity on the second lap. As shown above, the induction protocol in familiar environments has to be applied over several laps to successfully induce place field shift. In novel environments conversely, applying the induction protocol over a few laps is enough to induce the development of a new place field. Indeed, the induction of dendritic activity over 1-2 laps is sufficient to shift place field location (figure 5I-J). As initially hypothesized, our model indicates that we need fewer induction laps to induce place field shift in novel environments than in familiar ones. This extra plasticity of place fields in novel environments is due to two factors: synaptic weights are not yet strongly tuned in the first laps, and the novelty signal induces an increase in postsynaptic dendritic activity.

Finally, we investigate whether we can artificially manipulate the interneuron activity in familiar environments so that the model behaves like in novel environments. In particular, place fields could become more plastic. To test that, we run the simulations for 200 laps, thus until the environment becomes familiar. At lap 200, we decrease dendrite-targeting inhibition and increase soma-targeting inhibition, resetting them to the level of novel environment exploration. Simultaneously, at lap 200, we induce dendritic activity within a region far from the peak of the neuron’s place field. Since the modulation of inhibition is applied over the entire environment, there is an increase in both within- field and out-of-field firing rate. Accordingly, the shift in place field location is harder than in the case without manipulation of inhibition (figure 5K). We conclude that, surprisingly, resetting inhibition to novel environment levels is not enough to make place fields plastic again. Indeed, overall manipulation of inhibition reinforces stable place fields by increasing within-field activity.

In summary, our model suggests that single-cell place fields can be shifted under the induction of dendritic activity. Our model predicts that small induction areas are more efficient to induce the development of new place fields. Induction in novel environments is also more efficient than in familiar ones. Counter-intuitively, resetting novel environment level of inhibition represses place field plasticity.

## 3 Discussion

We propose a model of the hippocampal CA1 place cells in which interneuron activity is modulated by novelty in an interneuron-type-dependent manner. Using our simulations, we identify the potential mechanisms underlying the evolution of place fields and the transition from silent to place cells in novel environments. During the initial stages of exploration of novel environments, dendrite-targeting inhibition is reduced whereas soma-targeting inhibition is increased. The reduction in dendritic inhibition opens a window for plasticity, leading to the formation and stabilization of receptive fields. We then show that place fields are more stable in familiar environments than in novel environments. Our simulations suggest that this extra stability is due to stronger synaptic weights and increased dendritic inhibition. Our model makes predictions on how to perturb place fields by dendritic activation. In our model, dendritic activation can shift place field location. We predict that this shift is easier if the dendritic activity is induced only within a small region of the environment. We also predict that it is easier to induce place field shift in novel than in familiar environments. Our model, albeit simple, provides a mechanism for several features of the CA1 network and provides testable predictions.

The modulation of interneuron activity during exploration of novel environments is thought to be important for place field development and stabilization. Dendritic events, such as NMDA spikes and Ca^2+^ plateau potentials [Sheffield and Dombeck, 2015, Sheffield et al., 2017, Bittner et al., 2017], have been implicated in the development of new place fields. Thus, the reduction in dendritic activity might be responsible for opening a window for plasticity by promoting these dendritic events. Reduced inhibition could unmask small input inhomogeneities, leading to the rapid emergence of place cells during the first stages of exploration of novel environments. These small inhomogeneities would then be amplified through synaptic plasticity. The role of increased somatic inhibition in novel environments, however, is less clear. Since ST interneurons receive inputs from local pyramidal cells, the increase in ST interneuron activity could be reflecting the increase in pyramidal cell activity. Somatic inhibition can also be responsible for regulating pyramidal cell activity to ensure that the overall level of excitatory activity is kept within a certain regime. That might be important to ensure the development of sparse representations. This regulation could also be important to separate the learning process into two stages such that spatial representations are first developed within the hippocampus before being communicated back to the cortex. Furthermore, the increase in ST interneuron activity can also be responsible for controlling plasticity at CA1 pyramidal neurons.

Place cell firing rate has been shown to increase rapidly following exposure to novel environments [Frank et al., 2004, Cohen et al., 2017]. As suggested by Cohen et al. [Cohen et al., 2017], this increase is associated with increased excitatory inputs onto CA1 pyramidal cells in our model. Through exploration, pyramidal cell firing rate returns to baseline levels in familiar environments.

This later reduction in place cell firing rate has been suggested to be associated with a reduction in excitatory input [Cohen et al., 2017]. Conversely, inspired by data from Sheffield et al. [Sheffield et al., 2017], our model suggests that the return to baseline firing rate might be associated with the combined increase in dendritic inhibition and decrease in somatic inhibition, while excitatory inputs remain strong.

Our model indicates that the balance between strong excitatory input and the interplay between dendritic and somatic inhibition leads to place field stabilization in familiar environments. Synaptic plasticity, in our model, leads to the strengthening of within-field inputs and weakening of out-of- field inputs. The importance of plasticity for place field stabilization is corroborated by experiments in which NMDA receptors have been shown to be important for place field stabilization [McHugh et al., 1996, Kentros et al., 1998, Cacucci et al., 2007]. The increase in dendritic inhibition in familiar environments in our model induces a reduction in dendritic events and, thus, a reduction in plasticity- induced changes in place fields. Additionally, the combination of weak out-of-field inputs and strong dendritic inhibition leads to higher robustness to noise. Overall, place fields stabilize following the exploration of novel environments, in agreement with experiments [Cohen et al., 2017]. Besides changes in excitatory and inhibitory inputs, dendritic nonlinearities might also contribute to place field development and stabilization. In presence of noise, our model indicates that dendritic nonlinearities are crucial for reliable place field development. Therefore, our model offers a possible mechanism for place field stabilization and highlights the importance of interneuron diversity and the balance between strong excitatory and inhibitory inputs for this stabilization.

Our model provides a mechanistic understanding of the CA1 network. It reproduces a variety of observations, such as the dynamics of place fields during the exploration of novel environments and in familiar environments. Furthermore, we demonstrate that place fields can be manipulated by artificial depolarization of CA1 pyramidal cells in our model.

## Supporting information

## 4 Acknowledgments

This work was supported by CAPES Foundation (process n. 99999.001758/2015-02), Wellcome Trust, Simons Foundation, NIH, BBSRC, and EPSRC.

## 6 Methods

### 6.1 Neuron model

We use two-compartment, rate-based neuron models. Each neuron is modeled as two compartments: one representing the soma and another representing the dendrites. The dendritic compartment’s activity, *r*_*dend*_, is determined by

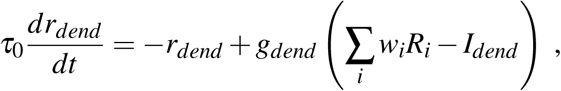

where *τ* _0_ is a time constant, *R*_*i*_ is the firing rate of neuron *i* in the presynaptic layer, *w*_*i*_ is the synaptic weight from a neuron in the presynaptic layer, *I*_*dend*_ is the input from dendrite-targeting interneurons— simulating SOM+ interneuron inputs—*g*_*dend*_ is a non-linear function of the input to the dendritic compartment given by

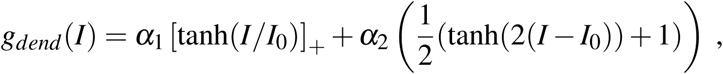

where [·]_+_ denotes a rectification that sets negative values to zero, *α* _1_ controls the linear gain of the dendritic compartment, *α* _2_controls the amplitude of the non-linear term associated with dendritic spikes, and *I*_0_ is proportional to the minimum input current necessary for the induction of a dendritic spike (supplementary figure 5A-B).

The somatic compartment receives input from the dendritic compartment and from soma-targeting interneurons—simulating PV+ interneuron inputs. The activity of the somatic compartment, *r*_*soma*_, is given by

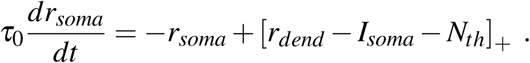

where *I*_*soma*_ is the input from soma-targeting interneurons—simulating PV+ interneuron inputs—and *N_th_* is the threshold for somatic activation.

### 6.2 Synaptic plasticity model

The synaptic weights from input neurons onto CA1 neurons are plastic and depends on the activity of the presynaptic neuron *r_j_*and the activity of the dendritic compartment of the postsynaptic neuron as a standard Hebbian term. We include a homeostatic term that takes into account the sum of all synaptic weights onto the postsynaptic neuron. The synaptic weight from input neuron *j* to the postsynaptic neuron *i*, *w_ij_* is updated following

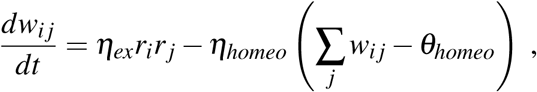

where *η_ex_* is the learning rate of excitatory connections, *η_homeo_* is the learning rate of the homeostatic term, and *θ_homeo_* is a target homeostatic constant.

### 6.3 Position-modulated inputs

The simulated CA1 neurons receive feedforward input from *N_pre_* neurons. These input neurons are tuned to specific locations and their firing rates span over the entire environment. All the place fields of input neurons have the same tuning width, *σ _pre_*, and the same amplitude, *A_pre_*. We assume that the animal explores an annular track of length *L* with speed *v*. The firing rate of an input neuron with place field centered at *p*_0_ is

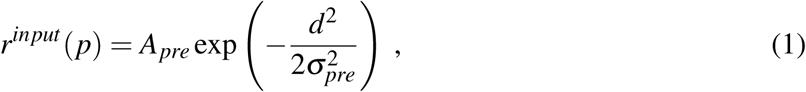

where *p* is the animal’s position, and *d* is the distance, along the track, between the animal’s position and the center of the place field.

### 6.4 Novelty signal

When simulating the exploration of a novel environment, we assume that the interneuron activity changes over time and is interneuron-type specific. We define a quantity, named novelty signal, that modulates the interneuron activity

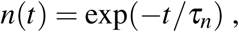

where *t* is the time measured from the start of exploration, and *τ _n_* is a time constant. The dendritic and somatic inhibition are then given by

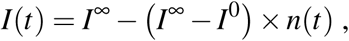

where *I^∞^* is the inhibitory activity in familiar environments, and *I*^0^ is the initial inhibitory activity in novel environments. The initial level of dendritic inhibition is assumed to be lower than its level in familiar environments, 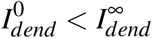 The initial level of somatic inhibition is assumed to be higher than its level in familiar environments,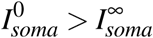.

### 6.5 Measuring place field stability

In section **??**, we analyze the stability of place fields in the first and last lap of novel environment exploration. In order to measure the effect of noise in novel environments, we go through the following steps: (1) we take the network in the state it was at the beginning of lap 1; (2) we simulate one lap of exploration, without plasticity; (3) we measure the place field of the postsynaptic neuron; (4) we rescale this place field such that its peak is set to 1; (5) we change the state of the network by adding noise to it (see below); (6) we repeat (2)-(4); (7) we calculate the absolute distance between the two rescaled receptive fields; (8) we repeat (6)-(7) *N_noise_* times and take an average over all samples (supplementary figure 5). To measure the effect of noise in familiar environments, we follow the same steps but using the state of the network at the beginning of the last lap (lap 50) in step (1).

We assume that place fields can be affected by three sources of noise: (i) noise at presynaptic place fields, (ii) noise at presynaptic firing rates, and (iii) noise at synaptic weights. In case (i), we multiply each presynaptic receptive field (equation 1) by a random variable taken from a normal distribution with mean 1 and variance *N*^2^. In case (ii), we assume that each presynaptic neuron receives an extra input, independent of its receptive field, and not tuned to the animal’s position. This extra input is taken from a normal distribution with mean 0 and variance *N*^2^ and then rectified to admit only positive values. In case (iii), we add a random number to each synaptic weight. This random number is taken from a normal distribution with mean 0 and variance *N*^2^. In all three cases, we define *N* as the noise amplitude.

### 6.6 Parameters and simulations

All simulations were implemented in python and are available at ModelDB. The parameters used in our simulations can be found in table 1

**Table 1:** Parameters summary

|  | Name | Value | Description |
| --- | --- | --- | --- |
| Neuron Model | $\tau_0$ | 2.0 ms | Firing rate time constant |
| | $\alpha_1$ | 2/3 | Linear gain of dendritic compartment |
| | $\alpha_2$ | 4/3 | Related to the amplitude of dendritic spikes |
| | $I_0$ | 5.0 | Minimum input to induce dendritic spikes |
| | $N_{th}$ | 1.0 | Threshold for somatic activation |
| Network Model | $N_E$ | 100 | Number of excitatory neurons in the network |
| | $N_I$ | 20 | Number of inhibitory neurons in the network |
| | $N_G$ | 10 | Number of groups of excitatory neurons |
| Plasticity Model | $\eta_{ex}$ | $5 \times 10^{-4} \text{ ms}^{-1}$ | Excitatory plasticity learning rate |
| | $\eta_{homeo}$ | $5 \times 10^{-5} \text{ ms}^{-1}$ | Homeostatic plasticity learning rate |
| | $\theta_{homeo}$ | 2.0 | Homeostatic target value |
| | $\eta_{in}$ | $1 \times 10^{-4} \text{ ms}^{-1}$ | Inhibitory plasticity learning rate |
| | $I_0^{ex}$ | 8.0 | Target excitatory input (for inhibitory plasticity) |
| | $A_{inhib}$ | 1.0 | Strength of inhibitory plasticity |
| | $A_{inhib} \text{ (S3H)}$ | 10.0 | Strength of inhibitory plasticity for supp fig 3H |
| Input | $A_{pre}$ | 2.0 | Presynaptic place field amplitude |
| | $\sigma_{pre}$ | 5.0 | Presynaptic place field width |
| Novelty signal | $\tau_n$ | $1.5 \times 10^{-4} \text{ ms}^{-1}$ | Time constant for novelty signal decay |
| | $I_{dend}^0$ | 0.0 | Initial dendritic inhibition |
| | $I_{dend}^\infty$ | 10.0 | Target dendritic inhibition |
| | $I_{soma}^0$ | 4.0 | Initial somatic inhibition |
| | $I_{soma}^\infty$ | 0.0 | Target somatic inhibition |

