## Supplementary material for "Interplay between somatic and dendritic inhibition promotes the emergence and stabilization of place fields"

### Supplementary Figures

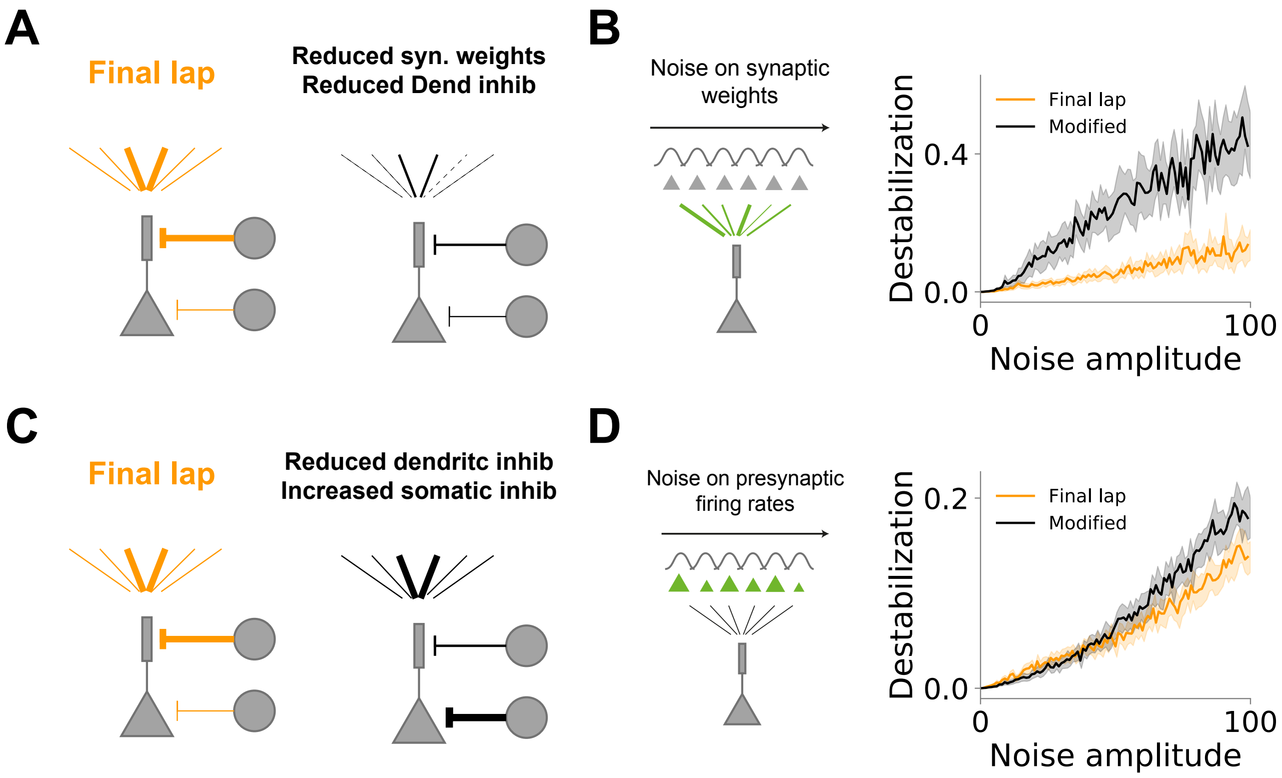


**Supplementary Figure 1 (related to figure 4). Strong synaptic weights and stronger dendritic inhibition ensures place field stability. (A-B)** Strong synaptic weights provide stability to noise on synaptic connections. **(A)** Left: Network diagram for the network state at the last lap of exploration in figure 4. Right: Modified network with reduced synaptic weights and reduced dendritic inhibition. Importantly, the changes are determined such that the neuron's place field is kept unchanged. **(B)** Destabilization of place fields by noise on synaptic weights for final lap of exploration (orange) and modified network as in (B) (black). **(C)** Left: Network diagram for the network state at the last lap of exploration in figure 4. Right: Modified network with reduced dendritic inhibition and increased somatic inhibition. Importantly, the changes are determined such that the neuron's place field is kept unchanged. **(D)** Destabilization of place fields by noise on presynaptic firing rates for final lap of exploration (orange) and modified network as in (C) (black).

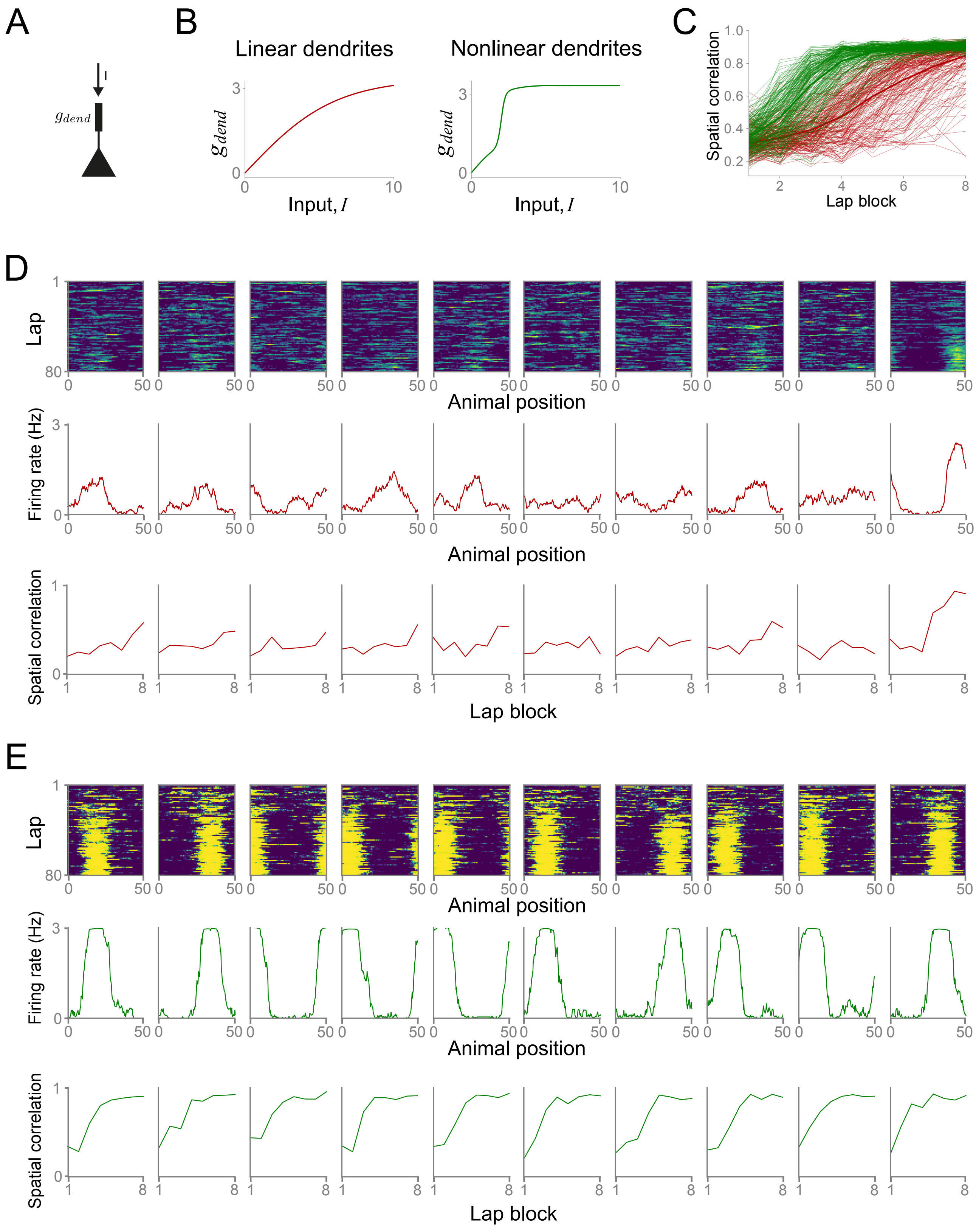
A
B
C
D
E
I
$g_{dend}$
Linear dendrites
Nonlinear dendrites
Input, $I$
Spatial correlation
Lap block
Lap
Animal position
Firing rate (Hz)

**Supplementary Figure 2. Dendritic non-linearity leads to reliable place field development.** **(A)** Single-cell diagram. A pyramidal neuron receives input $I$ and integrates it through a function $g_{dend}$. **(B)** Dendritic transformation function $g_{dend}$ as a function of the input $I$ for linear dendrite (left, red) and nonlinear dendrites (right, green). **(C)** Spatial correlation between laps for blocks of 10 laps on simulations with nonlinear dendrites (green) and linear dendrites (red). Thick lines show averages over 200 cells for each group. Thin lines are individual cells. Note that the spatial correlation for several cells with linear dendrites does not increase over lap blocks. **(D)** Examples of individual pyramidal cells with linear dendrites. Top, evolution of neuron firing rate over laps as a function of the animal position. Middle, average neuron firing rate over the last 10 laps of exploration as a function of the animal position. Spatial correlation between laps for blocks of 10 laps. **(E)** Examples of individual pyramidal cells with nonlinear dendrites. Top, evolution of neuron firing rate over laps as a function of the animal position. Middle, average neuron firing rate over the last 10 laps of exploration as a function of the animal position. Spatial correlation between laps for blocks of 10 laps.

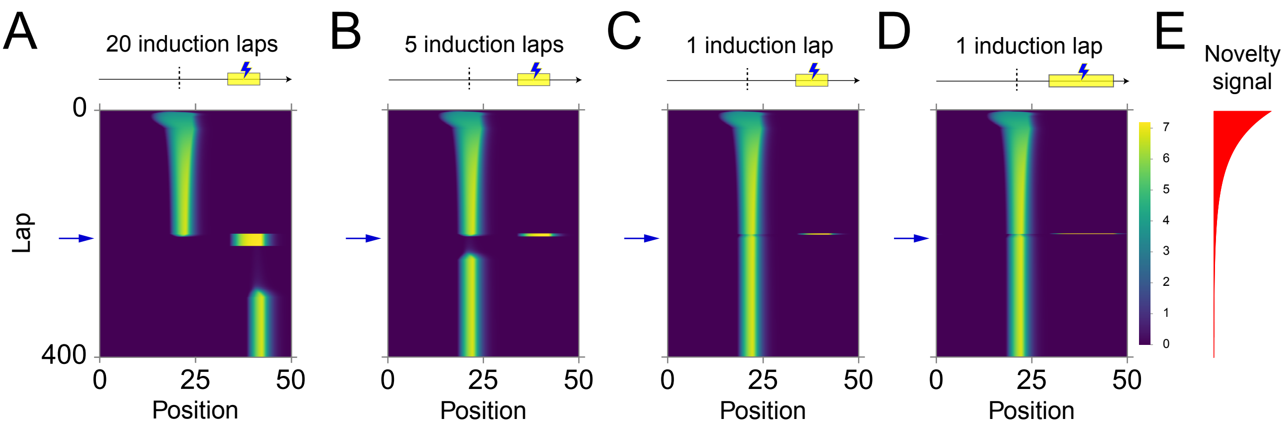


**Supplementary Figure 3 (related to figure 5). Artificially induced CA1 single cell activity can shift place field location. (A-D)** Evolution of place fields for the case in which an extra current is applied to the postsynaptic neuron while the animal traverses a section of the track. Yellow bar indicates the induction region in which the extra current is applied. Dashed line indicates the position of the peak of the initial place field. Blue arrow indicates the first induction lap (lap 200). **(A)** Place field evolution for 20 induction laps and small (1/6 of the track) induction region. Place fields are shifted towards new position determined by the region of extra current application. **(B)** Place field evolution for 5 induction laps and small (1/6 of the track) induction region. Place fields are transiently removed by the application of extra current and reemerge in the initial location. **(C)** Place field evolution for 1 induction lap and small (1/6 of the track) induction region. Place fields are not disturbed following the application of extra current. **(D)** Place field evolution for 1 induction lap and large (1/3 of the track) induction region. Place fields are not disturbed following the application of extra current. **(E)** Evolution of novelty signal over laps for the simulations in (A)-(D).

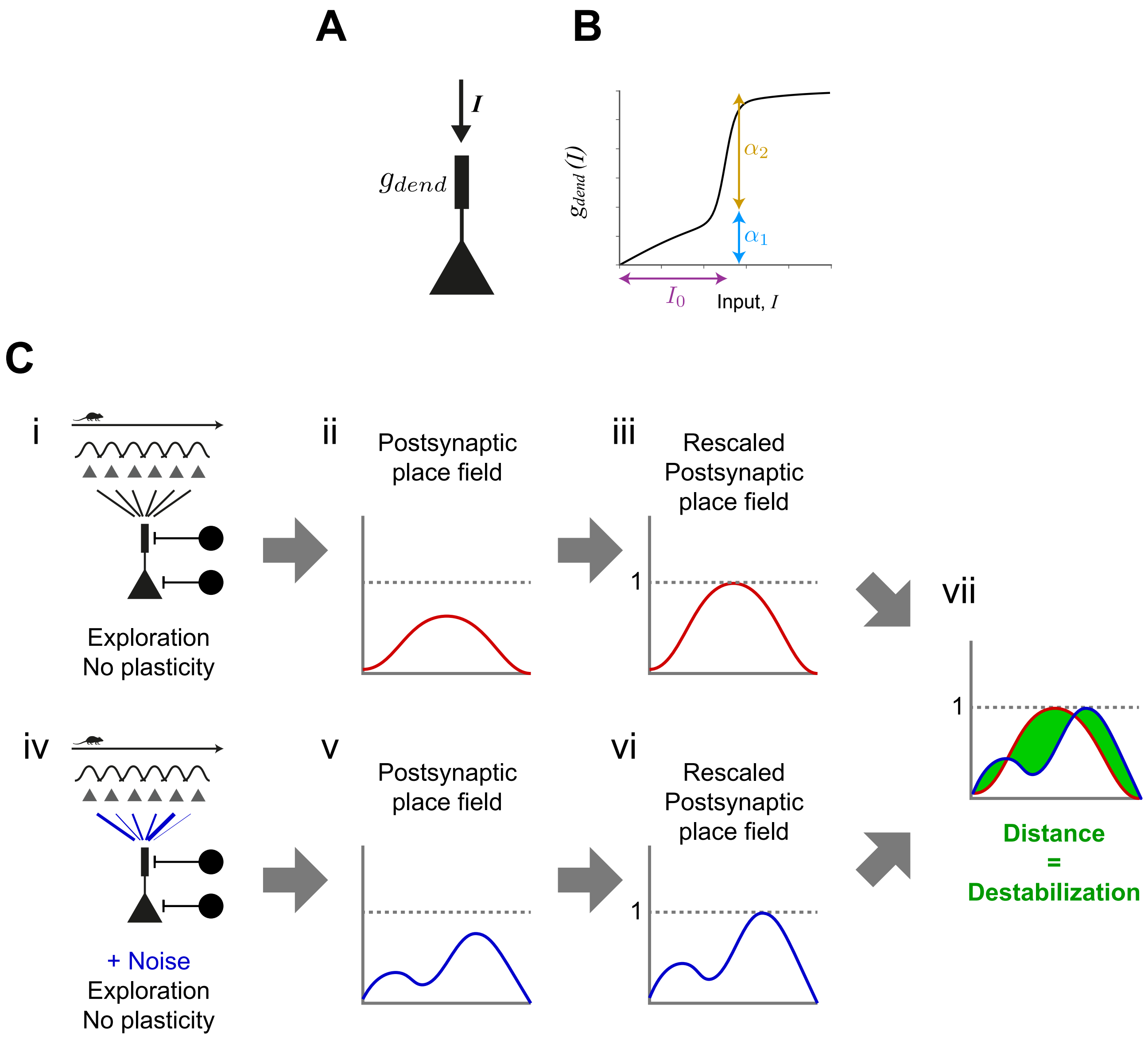


**Supplementary Figure 4. Dendritic non-linearity and stability analysis procedure.** **(A)** Single-cell diagram. A pyramidal neuron receives input $I$ and integrates it through a function $g_{dend}$. **(B)** Diagram of $g_{dend}$ as a function of the input $I$ (see methods). $\alpha_1$ controls the linear gain of the dendritic compartment; $\alpha_2$ controls the amplitude of the non-linear term related to dendritic spikes; and $I_0$ controls the minimum input to elicit dendritic spikes. **(C)** Place field stability analysis. For each measurement of place field stability (see methods) we perform the following steps: (i) we simulate one lap of exploration, without plasticity; (ii) we measure the place field of the postsynaptic neuron; (iii) we rescale this place field such that its peak is set to 1; (iv) we change the state of the network by adding noise to it; (v-vi) we repeat (ii)-(iii); (vii) we calculate the absolute distance between the two rescaled receptive fields.
